# Social isolation biases female rats toward safety-oriented, efficient foraging

**DOI:** 10.64898/2026.07.03.736107

**Authors:** Yadong Dai, Alexandra K. L. Seielstad, James R. Hinman

## Abstract

Social isolation has profound effects on behavior and cognition, but these effects can differ across sex and behavioral domains. We asked how social isolation shapes foraging decisions in male and female rats using a patch-leaving task that combines spatial navigation with sequential stay-or-go choices across varying travel costs and reward depletion rates. All groups scaled patch residence times with travel cost in a manner consistent with the marginal value theorem yet consistently overstayed beyond the optimal leaving time. The magnitude of overstaying was shaped by a strong interaction between sex and social status, with socially isolated females leaving patches closest to optimal and consuming food at the highest rate. Their foraging was accompanied by a coherent spatial and behavioral profile where the socially isolated females spent more time idle, preferentially occupying protected regions near the door and corridor, and avoiding the exposed patch center during both active foraging and periods of idling. These patterns are consistent with a conservative, safety-oriented strategy that simultaneously minimizes exposure and maximizes caloric return. Social isolation does not uniformly impair cognition but can selectively bias female rats toward efficiency-maximizing foraging decisions, consistent with the ecological pressures faced by outcast females in wild rat colonies.

## Introduction

*Rattus norvegicus*, the Norway rat, is a highly social animal that lives in colonies of up to hundreds of individuals organized by a hierarchical social structure. Social rank shapes an individual’s access to resources, with higher-status animals gaining preferential access to food, water, and mating partners. Individuals at the bottom of the hierarchy may ultimately be expelled from the colony and forced to strike out on their own as socially isolated outcasts. Such outcasts must navigate and forage under the persistent risk of encountering colony members, a pressure expected to shape their decisions about when to explore and when to exploit a food patch. The foraging decisions of an outcast rat therefore unfold under fundamentally different social and ecological constraints than those of an integrated colony member.

Optimal foraging theory holds that animals make decisions that maximize their resource intake per unit of time. The marginal value theorem (MVT), one formalization of this principle, has proven broadly accurate in describing the patch-leaving decisions made by many species, including parasitoid insects,^1,2^ rodents,^3–11^ birds,^12–14^ non-human primates,^15–17^ and humans.^5,18–21^ MVT predicts the optimal moment at which an animal should abandon the currently harvested patch and travel to a new one, given the travel time between patches and the rate at which resources deplete within them.^22^ Although a wide range of species behave in qualitative agreement with MVT, increasing patch residence time as travel cost rises, virtually all of them overstay, remaining in patches beyond the optimal leaving time and departing with marginal intake rates below the environmental average. Overstaying is among the most reproducible departures from optimality in foraging behavior, yet how the magnitude of overstaying is shaped by sex, social experience, and developmental history remains largely unexamined.

Patch-leaving decisions also vary substantially among individuals, and one consistent axis of this variability is sex. Female rats tend to leave patches earlier than males and differ in their sensitivity to manipulations of travel cost and reward depletion,^9^ paralleling sex differences reported in adjacent decision domains such as impulsive choice and delay discounting.^23^ Foraging in open environments further requires animals to trade energy intake against exposure to risk. Rats have an innate fear of open spaces, owing to evolutionary pressures of predation, and preferentially occupy the protected perimeter of an arena rather than its exposed center, a thigmotaxic tendency long used as an index of anxiety-related behavior^24^, with females typically showing stronger center avoidance than males. Because time spent in protected regions trades against time available to harvest, the spatial organization of foraging effort and the patch-leaving decisions it produces are unlikely to be independent, yet the two are rarely characterized together within a single controlled paradigm.

Adolescence is a sensitive period for the maturation of prefrontal circuitry, the hypothalamic-pituitary-adrenal (HPA) axis, and social cognition.^25^ Post-weaning social isolation in rodents produces persistent, often developmentally specific changes that extend into adulthood, including deficits in learning, memory, and impulse control,^26,27^ increases in anxiety and stress reactivity,^28,29^ fragmented sleep structure,^30,31^ and a weakened immune system.^32,33^ Critically, these consequences are not uniform across sex: females show greater HPA-axis reactivity to isolation than males^28,29^ and stronger or qualitatively different effects on anxiety and spatial learning,^26,34,35^ whereas isolation effects on impulsivity and reward processing have been described primarily in males.^23,27^ Adolescent social isolation therefore provides a well-defined manipulation for asking how the early social environment shapes adult decision-making in a sex-dependent manner.

In the wild, socially isolated outcasts experience markedly different foraging pressures than integrated colony members. Outcast rats have been described as approaching and withdrawing from patches regardless of the presence of conspecifics, avoiding patches occupied by other rats altogether, and restricting themselves to the peripheral, safer regions of a foraging environment even in the absence of conspecifics, all while bearing elevated predation and aggression risk.^36^ These pressures predict that adolescent social isolation, by approximating several experiential features of outcast life, should bias foraging toward cautious, efficiency-maximizing strategies, and that this bias should be most pronounced in females. To test this, we used a sequential two-patch foraging task with manipulated travel costs and reward depletion rates in male and female Long-Evans rats reared from postnatal day 28 in either same-sex social groups or social isolation. We asked whether socially isolated rats deviate less from MVT-optimal leaving times than group-housed rats, whether any such effect is sex-dependent, and whether shifts in foraging efficiency are accompanied by corresponding changes in the spatial organization of foraging behavior.

## Results

### Patch-leaving foraging task

Given the differences between socially isolated and socially integrated wild rats in foraging behavior and impaired performance of socially isolated laboratory rats on most cognitive tasks, we utilized a laboratory-based patch-leaving decision-making task in male and female rats that were either socially integrated or socially isolated at postnatal day 28. Rats (n = 18 male, n = 18 female) were randomly assigned to either the socially isolated or socially integrated housing condition, resulting in four groups (solitary male (SM), solitary female (SF), group male (GM), and group female (GF)), with their housing condition continuing throughout the entirety of the experiment. After four weeks of living in their assigned housing condition, rats were individually tested in a sequential patch-leaving foraging task in an arena that consisted of two square open field patches connected by a corridor that allowed for the imposition of different travel times between the patches by restricting the rat to the corridor for different periods of time (Figure 1A). Following entry into a patch after the imposition of a travel time in the corridor, rats could forage for sucrose pellets distributed according to one of four different depletion functions that varied based on two initial pellet rates and two depletion rates (Figure S1). Rats readily shuttled between the two patches even with the imposition of different travel times (either 7 or 60 seconds) in the corridor, with the pellet distribution function resetting upon reentry into a patch after having visited the other patch.

**Figure 1.**
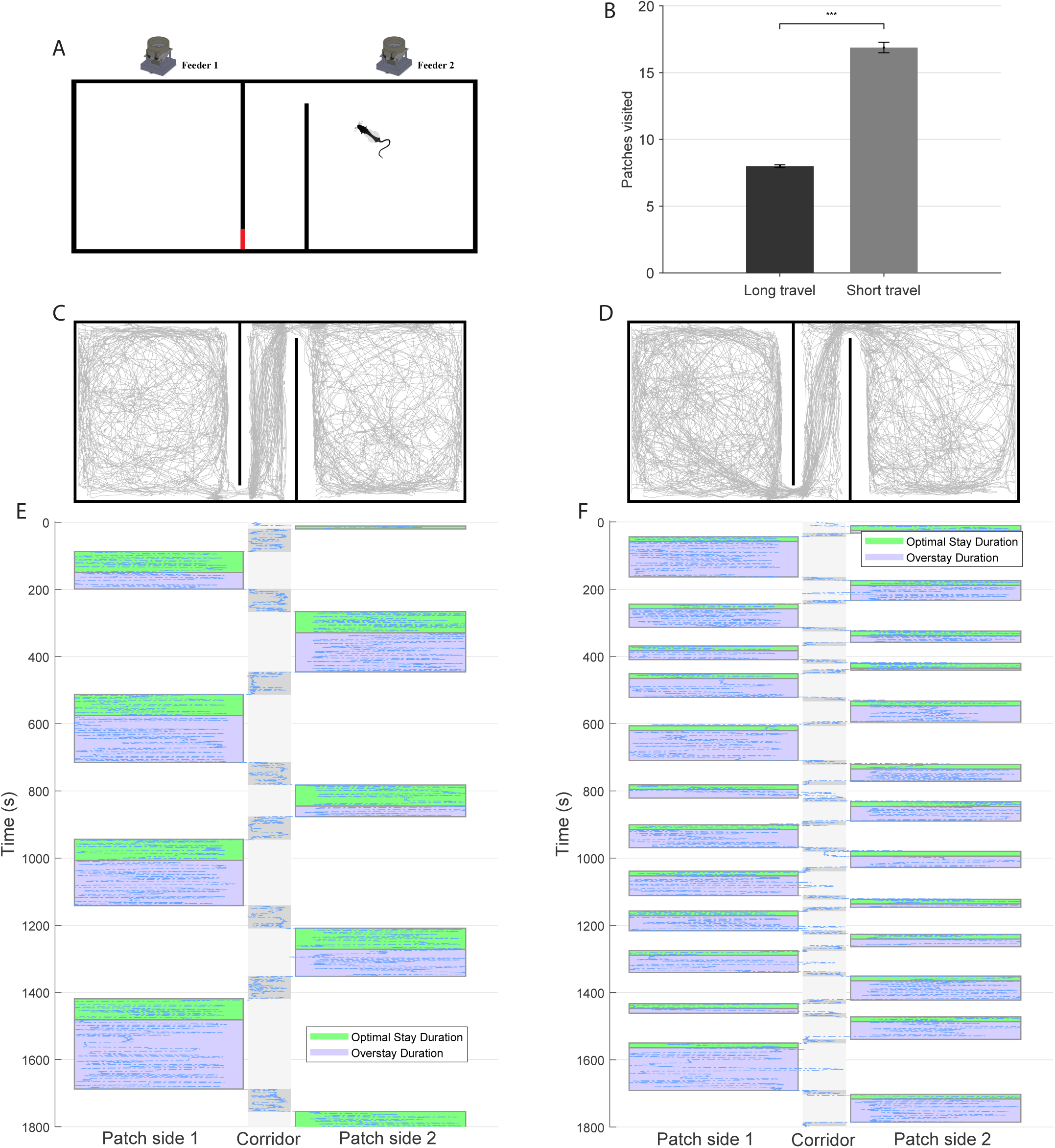
Rats adjust patch-leaving behavior according to travel cost in a sequential foraging task. (A) Schematic of the patch-leaving foraging arena. (B) Number of patches visited during long versus short travel time sessions. (C, D) Representative trajectory illustrating a rat’s movement pattern across the two patches and the corridor under long (C) and short (D) travel-cost conditions. (E, F) Example session activity plots for a long (E) and short (F) travel time session showing the rat’s trajectory in just the x-dimension over time (y-axis). Upon each entry into a patch the optimal leaving time is shaded in green, with any overstay duration shaded in blue.

Rats visited more patches with a short travel time (16.90+/-0.39 patches) than with a long travel time (8.00+/-0.11 patches; t(286)=21.9, p<0.001; Figure 1B). The marginal value theorem predicts that animals’ patch residence times will vary directly with the travel time between patches, which has been observed in both wild and laboratory animals, and was observed in this laboratory-based patch-leaving task (Figure 1C-F). Rats had significantly longer patch residence times with long versus short travel times (long: 174.53 ± 3.66 s; short: 102.92 ± 3.10 s; t(286) = 14.9, p < 0.001; Figure 2A). Rats’ patch residence times were compared to the optimal patch leaving time predicted by MVT, and as is typical for most animals, rats overstayed in patches beyond the optimal leaving time regardless of travel time or pellet distribution function, with rats overstaying significantly longer when there was a short as compared to a long travel time (long: 52.95 ± 5.16 s; short: 82.77 ± 3.11 s; t(286) = 4.95, p < 0.001; Figure 2B). The four pellet distribution functions did not result in differences in either the number of patches visited (high initial/low depletion: 11.6 ± 0.61; high initial/high depletion: 12.6 ± 0.64; low initial/low depletion: 12.7 ± 0.67; low initial/high depletion: 12.9 ± 0.72; one-way ANOVA: F(3,284) = 0.79, p = 0.499; Tukey–Kramer, all p > 0.05; Figure 2C) or patch residence times (high initial/low depletion: 143.9 ± 6.6 s; high initial/high depletion: 135.7 ± 6.1 s; low initial/low depletion: 135.7 ± 6.4 s; low initial/high depletion: 139.6 ± 6.5 s; one-way ANOVA: F(3,284) = 0.36, p = 0.779; Tukey–Kramer, all p > 0.05; Figure 2D). As rats were tested repeatedly across eight days with different pellet distribution functions and travel times, we considered whether rats’ memory of the previous session influenced their initial patch leaving decisions during the current session. Using the patch residence time of the first trial in each session, we found no differences across sessions (repeated-measures ANOVA, trials: F(7,238) = 1.35, p = 0.227), between sexes (sex × trials: F(7,238) = 0.42, p = 0.889; Figure 2E), or between social status (housing × trials: F(7,238) = 0.54, p = 0.807; 2F). This suggests that rats did not carry over specific patch-leaving biases from day to day.

**Figure 2.**
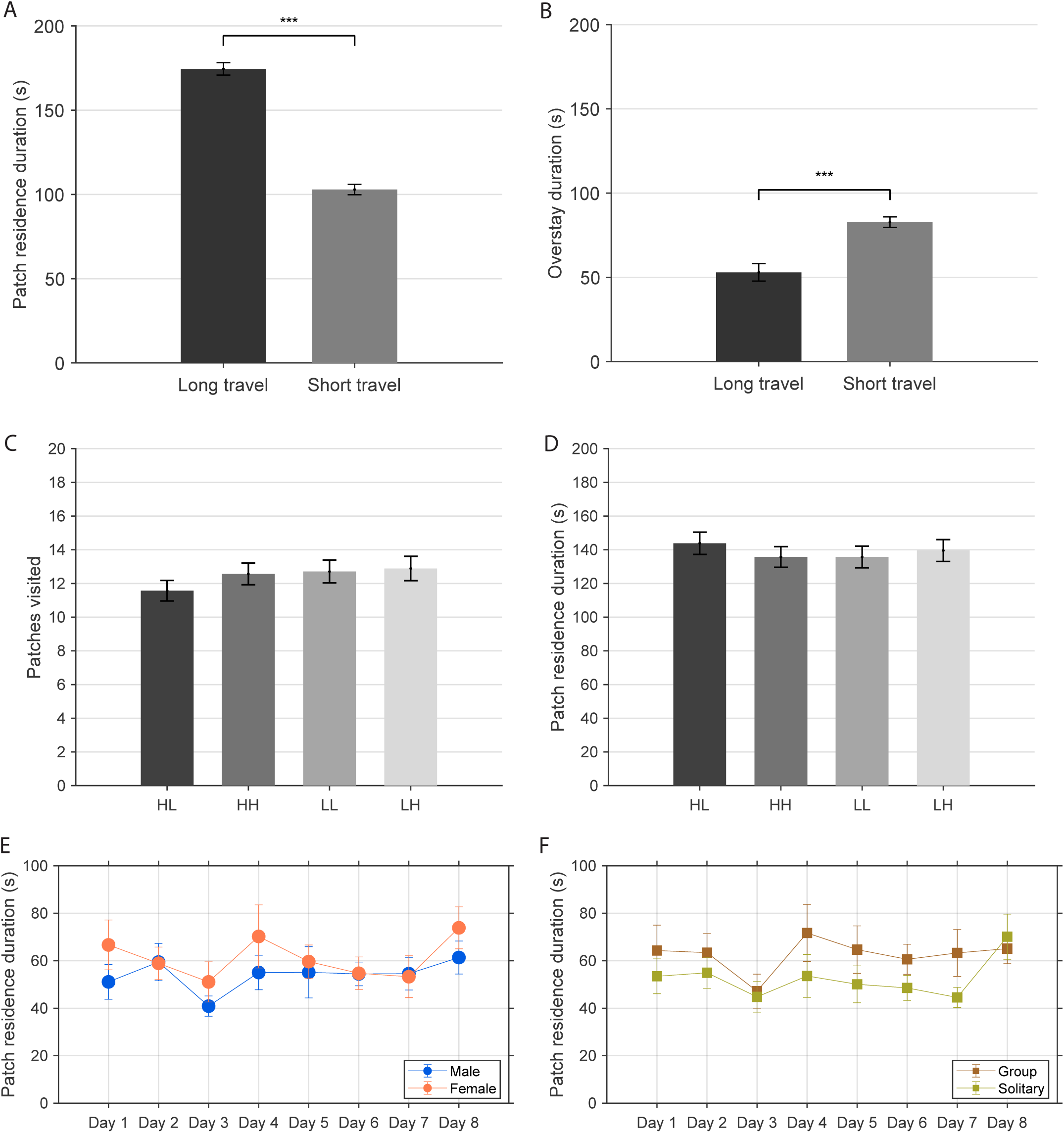
Patch residence times scale with travel cost and reflect systematic overstaying relative to marginal value theorem predictions. (A) Mean patch residence time for long and short travel time sessions. (B) Mean overstay duration for long and short travel time sessions. (C) Mean number of patches visited for the four different pellet distribution functions. (D) Mean patch residence times for the four different pellet distribution functions. (E) Mean patch residence times on the first trial of each session across days for males and females. (F) Mean patch residence times on the first trial of each session across days for group-housed and solitary rats.

### Socially isolated female rats make efficient foraging decisions

Each group of rats displayed overstaying behavior, yet clear differences were observed between the sexes and housing conditions. There is a strong interaction between sex and housing condition, driven primarily by socially isolated female rats, which overstay significantly less than the other groups (GM: 65.9 ± 4.2 s; SM: 77.8 ± 8.3 s; GF: 84.8 ± 6.1 s; SF: 46.9 ± 5.2 s; two-way ANOVA main effect of housing: F(1,32) = 4.44, p = 0.043; sex × housing: F(1,32) = 16.31, p < 0.001; Post hoc Tukey–Kramer: SF<SM: p<0.01, SF<GF: p<0.001; Figure 3A). Thus, socially isolated female rats perform closer to optimal in terms of their patch residence times. Given that an animal that leaves patches at the optimal time will maximize their food intake, socially isolated female rats consumed pellets at a significantly greater rate than each of the other three groups (GM: 4.44 ± 0.09; SM: 4.44 ± 0.16; GF: 4.40 ± 0.08; SF: 4.89 ± 0.10; two-way ANOVA main effect of housing: F(1,32) = 5.19, p < 0.05; main effect of sex: F(1,32) = 3.38, p = 0.075; sex × housing interaction: F(1,32) = 5.19, p < 0.05; Post hoc Tukey–Kramer: SF<GF, SM and GM all p<0.05; Figure 3B). This positions solitary female rats as more efficient foragers for food as a function of time than the other groups.

**Figure 3.**
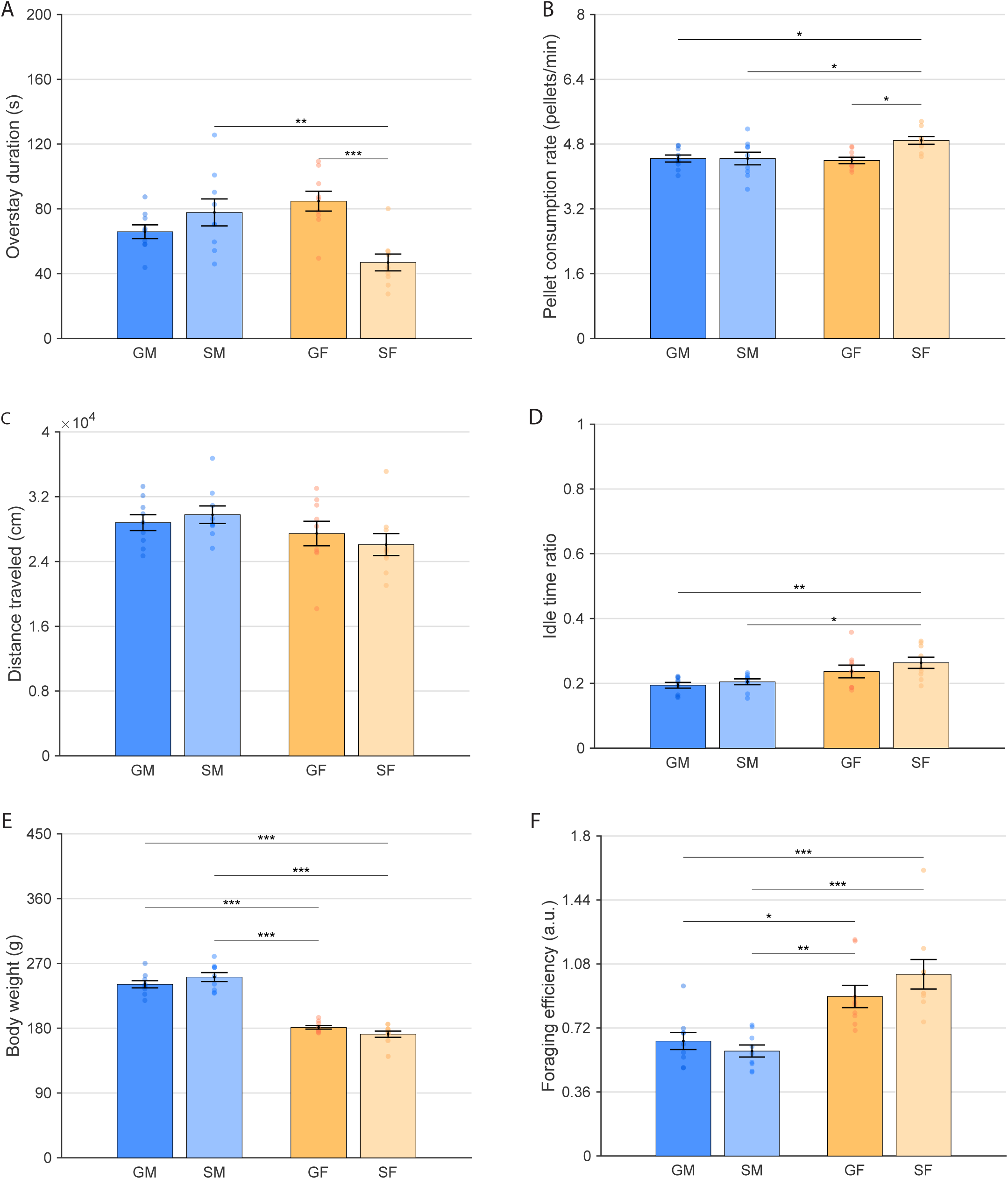
Social isolation selectively enhances foraging efficiency in female rats. (A) Mean patch overstay durations. (B) Mean pellet consumption rate. (C) Mean distance traveled. (D) Mean ratio of time spent idle. (E) Mean body weight. (F) Mean foraging efficiency.

As optimal foraging theories are ultimately concerned with animals making decisions to meet their caloric needs, an animal’s energy expenditure and overall caloric needs also play into whether animals are making optimal or efficient decisions. As the task structure does not require the rats to be constantly active, given that food pellets are dispensed into the patch according to the distribution function and not based on the rats’ behavior, the rats do not need to do anything during the inter-pellet interval and can simply choose to sit idly and conserve their energy. As such, on the energy-expenditure side, we considered (1) the total distance traveled during the patch-residence period and (2) the amount of time spent idle during the task as proxies for energy expenditure. There were no significant differences in travel distance during patch residence (GM: 28,803 ± 982 cm; SM: 29,781 ± 1,084 cm; GF: 27,457 ± 1,516 cm; SF: 26,086 ± 1,359 cm; two-way ANOVA main effect of housing: F(1,32) = 0.02, p = 0.876; main effect of sex: F(1,32) = 4.04, p = 0.053; sex × housing interaction (F(1,32) = 0.88, p = 0.356); Figure 3C), but solitary females spent significantly more time idle than both male groups (GM: 0.19 ± 0.01; SM: 0.20 ± 0.01; GF: 0.24 ± 0.02; SF: 0.26 ± 0.02; two-way ANOVA main effect of sex: F(1,32) = 12.18, p<0.001; main effect of housing: F(1,32) = 1.65, p = 0.209; sex × housing interaction: F(1,32) = 0.31, p = 0.580; post hoc Tukey–Kramer: SF>GM, p<0.01, SF>SM, p<0.05; Figure 3D). As female rats are significantly smaller than male rats (GM: 241.0 ± 5.0 g; SM: 251.1 ± 6.2 g; GF: 181.3 ± 2.6 g; SF: 171.7 ± 4.4 g; two-way ANOVA main effect of sex: F(1,32) = 217.82, p < 0.001; main effect of housing (F(1,32) = 0.00, p = 0.960; sex × housing interaction: F(1,32) = 4.32, p<0.05; post hoc Tukey–Kramer: males>females, all p < 0.001; Figure 3E), their foraging behavior makes them more efficient than at first glance, given their lower caloric needs. We derived a measure of foraging efficiency that incorporated the number of food pellets obtained, distance traveled, and body weight. Using this measure, female rats, particularly solitary females, were more efficient foragers than male rats (GM: 0.65 ± 0.05; SM: 0.59 ± 0.03; GF: 0.90 ± 0.06; SF: 1.02 ± 0.08; two-way ANOVA main effect of sex: F(1,32) = 32.67, p < 0.001; main effect of housing: F(1,32) = 0.33, p = 0.57; sex × housing interaction: F(1,32) = 2.29, p = 0.14; post hoc Tukey–Kramer: females>males, all p < 0.05; Figure 3F). Overall, these results indicate that solitary female rats were the most efficient foragers among the four groups tested, as they left patches earlier, obtained food pellets at a higher rate, and reduced locomotion during patch foraging relative to the other groups.

### Socially isolated female rats utilize space differently

Alternatively, reduced locomotor activity in solitary female rats may represent a safety-oriented foraging strategy that minimizes exposure and reduces the likelihood of encountering conspecifics. Rats’ overt utilization of space suggests differences in foraging strategies across the groups. Rats have an innate fear of open spaces owing to evolutionary pressures of predation occurring in such spaces, and therefore rats prefer the relative safety of protective structures, including the walls of laboratory-based arenas. Solitary, outcast wild rats have been described as restricting themselves to the perimeter of a foraging area,^36^ even in the absence of conspecifics, and therefore, we considered how rats utilize space in the laboratory-based foraging arena. During patch residence, rats in all groups preferentially occupied regions near the perimeter of the patches rather than the exposed central area (Figures 4A–4D). This tendency was more pronounced in females, which exhibited lower center-to-perimeter occupancy ratios than males, indicating reduced use of the patch center by female rats. Consistent with this pattern, male rats showed higher center: perimeter ratios than females, reflecting greater occupancy of the exposed central region during active foraging. In addition to the patch perimeter, the corridor and door regions constitute relatively protected areas of the environment. Because door closure and patch switching were triggered only when rats crossed the far end of the corridor, animals could remain near the door of the current patch without initiating the travel time to the other patch. Using the region definitions illustrated in Figure 4E, we quantified spatial occupancy across the full patch-residence period (Figure 4F). Significant group effects were observed in the Door, Corridor, and Patch Center regions, but not in the Boundary (one-way ANOVAs Door: F(3,284) = 8.06, p < 0.001; Corridor: F(3,284) = 16.47, p < 0.001; Patch Center: F(3,284) = 19.98, p < 0.001; Boundary: F(3,284) =<u>1.70,</u> p = 0.17). Solitary females spent a greater proportion of time near the door than both group-housed and solitary males (post hoc Tukey-Kramer Door: SF>GM p<0.01, SF>SM p<0.001), whereas both female groups spent more time in the corridor than males (post hoc Tukey-Kramer Corridor: SF>SM & GM p<0.001, GF>SM & GM p<0.001). Conversely, solitary females spent less time in the Patch Center than all other groups (Patch Center: SF<GF p<0.05, SF<SM p<0.001, SF<GM p<0.01; Figure 4F).

**Figure 4.**
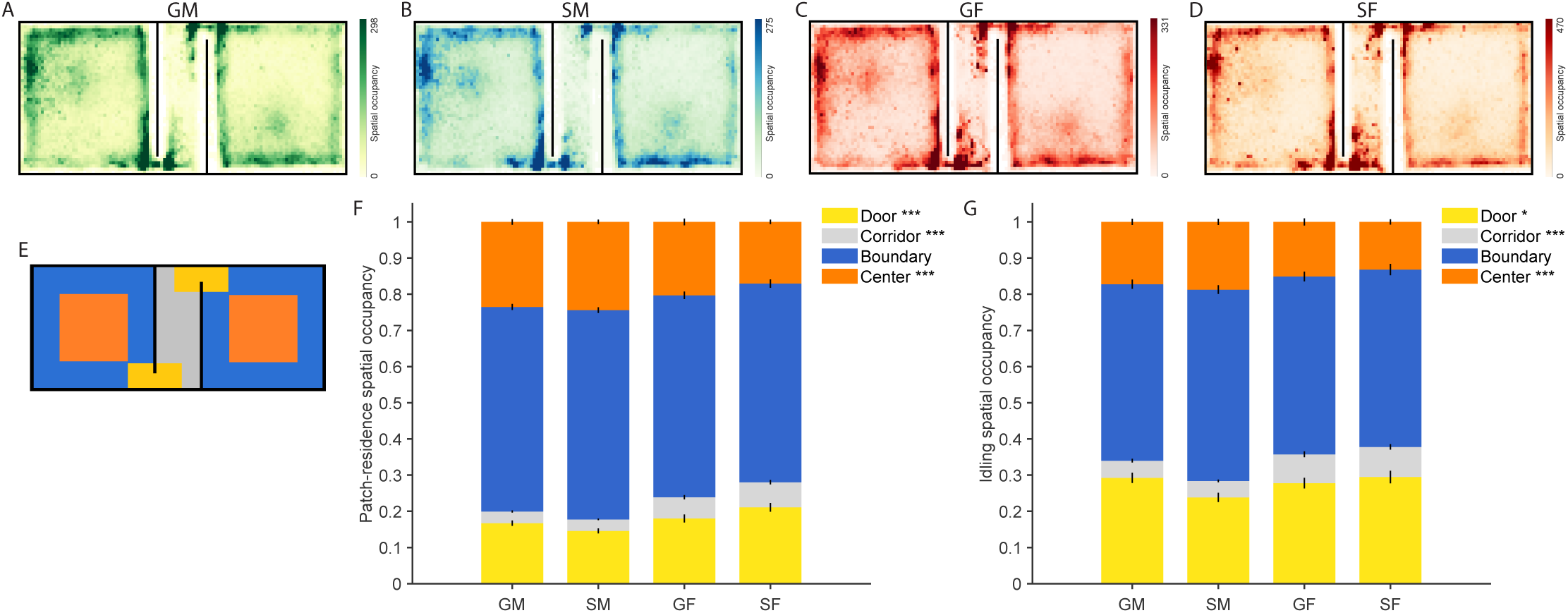
Social isolation and sex shape spatial occupancy during patch residence and idling. (A–D) Heat maps showing the mean spatial occupancy for group housed males (A), solitary males (B), group housed females (C), and solitary females (D). Color scale indicates relative occupancy, with armer colors denoting greater time spent in a given location. (E) Schematic of the foraging arena illustrating region definitions used for spatial analyses. Blue denotes the patch boundary (periphery), orange denotes the patch center, yellow denotes the area near the door of the current patch, and gray denotes the corridor connecting the two patches. (F) Proportion of spatial occupancy in the Door, Corridor, Boundary, and Patch Center regions during patch residence for group housed males, solitary males, group housed females, and solitary female group. (G) Proportion of spatial occupancy in the Door, Corridor, Boundary, and Patch Center regions during idle time for group housed males, solitary males, group housed females, and solitary female group.

To determine whether these spatial biases persisted during non-foraging periods, we next examined spatial occupancy during idling (Figures 4G). Group differences were again observed in the Door, Corridor, and Patch Center regions, but not in the Boundary (one-way ANOVAs Door: F(3,284) = 3.12, p < 0.05; Corridor: F(3,284) = 10.08, p < 0.001; Patch Center: F(3,284) = 8.15, p < 0.001; Boundary: F(3.284) = 2.09, p = 0.10). Solitary females spent more idle time in the door and corridor regions than solitary males (post hoc Tukey-Kramer Door: p<0.05, Corridor: p<0.001, whereas both female groups showed reduced occupancy of the Patch Center relative to males (post hoc Tukey-Kramer: SF<SM p<0.001, SF<GM, p<0.01, GF<SM p<0.05; Figure 4G). Together, these results indicate that solitary female rats preferentially restrict both foraging and idle behavior to regions associated with greater environmental safety, concentrating spatial use near the door and corridor while minimizing exposure to the open field.

## Discussion

The present study reveals a striking and counterintuitive pattern: social isolation, which typically impairs cognitive function across a host of laboratory-based tasks, was associated with more efficient foraging specifically in female rats. Socially isolated females left patches closer to the optimal time predicted by MVT than the other three groups, overstaying significantly less and acquiring sucrose pellets at a higher rate across both short and long travel costs. This shift in decision-making was accompanied by a coherent spatial and behavioral profile: solitary females preferentially occupied the protected regions near the door and corridor, avoided the exposed patch center during both active foraging and idling, and spent a greater fraction of the session idle. The convergence of efficiency-relevant decisions with safety-oriented spatial use indicates that adolescent isolation does not produce a domain-general impairment in female rats, but instead biases them toward a coordinated strategy that simultaneously minimizes exposure and maximizes caloric return—a strategy consistent with the ecological pressures faced by outcast females at the margins of wild rat colonies, where both predation risk and conspecific aggression are elevated.

The interaction between sex and social condition, driven largely by solitary females showing the lowest levels of overstaying relative to MVT, suggests that females may be more sensitive than males to the effects of social isolation, but that this sensitivity manifests in complex, domain-specific ways. The same manipulation that shifted female foraging toward a more conservative, efficiency-maximizing strategy impairs females more severely than males in other cognitive domains, including spatial learning and memory.^34,35^ This sex-specific pattern may have hormonal underpinnings. Post-weaning social isolation increases stress and anxiety sensitivity more strongly in females, which show greater HPA-axis reactivity following isolation,^28,29^ and ovarian hormones modulate anxiety-like behavior and stress responses across the estrous cycle,^37,38^ providing a plausible mechanistic link between social isolation, heightened anxiety, and altered foraging strategy. Because estrous-cycle phase was not monitored in the present design, a cycle-dependent contribution to the female-specific effects cannot be evaluated here; future studies should examine whether patch-leaving decisions vary across the estrous cycle or following ovariectomy.

The spatial restriction observed in solitary females extends Calhoun’s classical descriptions of outcast wild rats into a controlled laboratory paradigm.^36^ Across both active foraging and idle periods, solitary females concentrated their occupancy in the door and corridor regions and avoided the patch center, whereas group-housed females and both male groups distributed their occupancy more evenly. The persistence of this bias during idling is informative: a spatial preference expressed only during active foraging could reflect a tactical response to the depleting reward schedule, whereas one that holds during periods of disengagement points to a more generalized defensive stance. Center avoidance in open-field assays has long been interpreted as an index of state and trait anxiety^24^, and the present spatial pattern is consistent with that interpretation. The novel observation is that this defensive spatial profile coexists with, and likely directly supports, efficiency-maximizing patch-leaving decisions, rather than trading off against them.

Because the task delivered pellets on a fixed schedule rather than in response to behavior, rats did not need to remain active during the inter-pellet intervals and could conserve energy by sitting idly. The elevated idling of solitary females, therefore, admits two competing interpretations. Under a motivational-deficit account, isolation reduces engagement with the task, and the increased idle time reflects disengagement rather than strategy. Under an energy-conservation account, idling is a deliberate behavioral choice that lowers caloric expenditure during inter-pellet intervals and contributes to the observed foraging efficiency. Three features of the data favor the strategic interpretation. First, idling co-occurred with higher pellet-acquisition rates and closer-to-optimal patch leaving, rather than with reduced engagement. Second, idling was concentrated in the protected door and corridor regions rather than distributed uniformly, the spatial signature of an active safety-related preference rather than a global loss of behavioral drive. Third, solitary females continued to traverse the corridor and re-enter patches at appropriate intervals, indicating that the goal-directed structure of the task remained intact. A motivational-deficit account predicts disengagement that scales with idling, which neither the temporal nor the spatial structure of the present data supports.

Several mechanisms could account for the reduced overstay observed in solitary female rats. Overstaying in patch-leaving paradigms has been attributed to the undervaluation of the opportunity cost of remaining in the current patch,^19^ rational learning of patch structure under uncertainty,^21^ and choice-history biases that prolong patch residence.^5^ Adolescent social isolation could shift any of these computations, for example, by heightening the subjective salience of the opportunity cost, by altering prefrontal contributions to structure learning,^26,34,35^ or by changing subjective time perception under chronic stress. Notably, the direction of the effect argues against a generalized impulsivity account: testosterone has been implicated in sex differences in impulsivity, and removal of endogenous testosterone via orchiectomy increases impulsivity in males,^23^ so a reward-driven impulsivity mechanism would predict earlier patch leaving in males rather than the female-specific reduction in overstay observed here. Distinguishing among the remaining accounts will require within-task manipulations not implemented in this study, including reward-rate volatility, explicit cuing of patch structure, and pharmacological challenge of HPA-axis function.

Together, these findings contribute to a growing body of evidence that early social experience has profound and long-lasting effects on behavior, but that these effects are not uniformly detrimental across all domains. While social isolation impairs many forms of cognitive performance,^26,27,34,35^ it appears to enhance certain aspects of decision-making in the context of foraging, particularly for females and in situations where conservative, efficient strategies are advantageous. Consistent with re-wilding arguments that the magnitude and direction of social-adversity effects are strongly context-dependent in richer, more naturalistic social ecologies,^39^ our results suggest that adolescent social isolation can bias females toward an adaptive, efficiency-maximizing foraging strategy in specific contexts, such as patch-leaving. This context-dependence emphasizes the importance of evaluating the cognitive consequences of early-life social manipulations across a broad range of ecologically valid paradigms before characterizing them as global deficits, and of recognizing that behaviors appearing maladaptive in one context (e.g. reduced exploration, elevated thigmotaxis, increased idling) may be adaptive in another.

### Limitations of the Study

Several limitations of this study should be noted. First, the sample size was modest (n = 9 per group), although in line with other studies in the field; while the central sex × housing interaction on overstay duration and pellet-acquisition rate was robust, the body-weight-normalized foraging-efficiency metric showed a main effect of sex without a significant sex × housing interaction (p = 0.14), so the efficiency conclusion rests on the convergence of multiple measures rather than on a single significant interaction in the composite metric. Second, estrous-cycle phase was not monitored, which precludes direct evaluation of cycle-dependent contributions to the female-specific effects reported here. Third, isolation onset at postnatal day 28 captures a single developmental window, and the effects of isolation initiated at earlier or later ages may differ. Fourth, behavioral characterization was restricted to the patch-leaving paradigm; how the observed strategy generalizes to other foraging structures, to social foraging contexts, or to overtly threatening environments remains to be established. Finally, the present study is purely behavioral, and the neural substrates that support the observed sex × isolation interaction, including candidate contributions from anterior cingulate cortex,^8^ locus coeruleus,^6^ and HPA-axis circuitry, remain a priority for future investigation.

## STAR★METHODS

### KEY RESOURCES TABLE

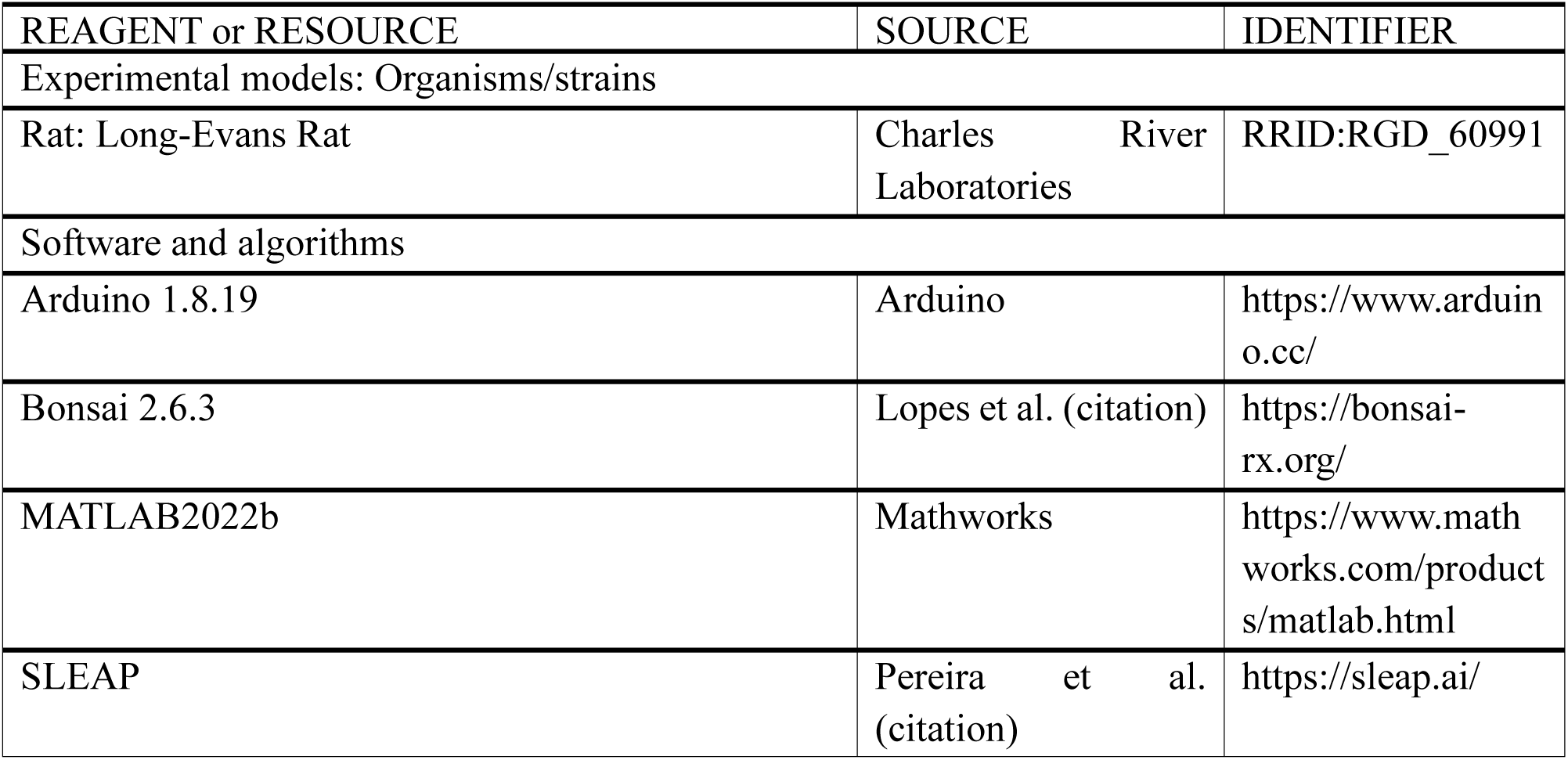

### EXPERIMENTAL MODEL AND SUBJECT DETAILS

The experimental protocols were approved by the Institutional Animal Care & Use Committee (IACUC) of the University of Illinois Urbana-Champaign. Thirty-six (18 males and 18 females) Long-Evans hooded rats (weight: 75 g, approximately 4 weeks old) were obtained from Charles River Laboratories in three rounds, each batch with 12 rats (6 males and 6 females). Animals were evenly assigned to housing either individually or in same-sex groups of three in standard polypropylene shoebox cages for four weeks, with water and food ad libitum for 4 weeks until task acclimation began. The room was maintained at a constant temperature of 23°C and on a 12-hour standard light/dark cycle. During training and testing sequences, animals’ body weights were maintained at >85% of their free-feeding weight to encourage food foraging.

### METHOD DETAILS

#### Behavioral apparatus

The behavioral apparatus was an integrated rectangular structure of gray acrylic panels, featuring two open fields (75 cm squares) connected by a corridor (75 cm x 25 cm). The walls of the apparatus were 45 cm tall. Extra maze cues were strategically placed around the maze to facilitate animals’ orientation. Access to each side of the open field was controlled by an automatic door with a stepper motor. Infrared light break beam sensors monitored if rats were sufficiently close to the automatic door and instructed its operations. A 3D-printed sugar pellet dispenser with a stepper motor was positioned above each open field. The dropped sugar pellets were scattered randomly across various locations in the open field. All stepper motors and infrared light break beam sensors were operated by an Arduino UNO board. Arduino codes and data were written and streamed through Arduino IDE 1.8.19. A FLIR Flea3 camera was placed above the apparatus to record animal behaviors. Behavioral data and video data were integrated and stored by Bonsai (an open-source software) on the PC platform.

#### Patch-leaving training and testing

After four weeks of separate housing conditions, rats underwent 5 days of handling and apparatus acclimation, with each handling session lasting 5 minutes. Following handling, the experimenter placed the rat in the corridor, with both doors on either side remaining open. In each of the open fields, 20 sugar pellets were pre-positioned. The rats were allowed to freely navigate between the patches for 10 minutes to retrieve the sugar pellets. During the 5-day training phase, we chose a short corridor transit time of 7 seconds and medium initial and depletion rates as the reward schedule. During the training sessions, the rats were placed in the corridor with both doors closed. After 10 seconds, one side door would randomly open, initiating a new reward schedule in the corresponding open field, the rat was allowed to stay in that patch for any length of time. When a rat chose to leave the patch and entered the corridor, approaching the opposite automatic door and triggering the infrared light break beam sensor, the door of the previous patch closed. After the transit time elapsed, the door on the other side opened, starting a new reward schedule in the new patch. Rats could repeatedly navigate between the two open fields within 30 minutes. Throughout the 5-day training procedure, we consistently used this reward rate. The testing phase lasted for 8 days. The experimental procedure was identical to the training phase, with the difference being that during these eight days, we employed different transit times (short-7 seconds and long-60 seconds), initial rates (low and high), and depletion rates (low and high). This resulted in 8 different combinations (2^3) of parameters. Over eight days, each rat experienced the eight reward schedules, each session lasting 30 minutes. These reward schedules were pseudo-randomly ordered, and all three batches of rats underwent the same sequence of reward schedules. Rats underwent handling, acclimation, training, and testing individually. Housing conditions remained constant as separate housing conditions throughout this period.

#### Reward schedules

The reward schedule was determined by three parameters: corridor transit time, initial rate, and depletion rate. We used dustless precision pellets (F0023, 100% sucrose, 45 mg, Unflavored) as rewards. Corridor transit time was defined as the waiting time in the corridor when both doors are closed, occurring between a rat abandoning the previous patch and moving to the next one. We selected two corridor transit times: 7 seconds and 60 seconds. The initial rate represented the initial interval between sugar pellet dispensing, with a low initial rate of 20 seconds and a high initial rate of 5 seconds. The depletion rate was the weighted interval time added to the initial interval time. A higher depletion rate resulted in a faster increase in the interval time between sugar pellet dispensing as the cumulative residence time lengthened. Conversely, a lower depletion rate led to smaller changes in the interval time between sugar pellet dispensing. Together, the initial rate and depletion rate determined the interval schedules for sugar pellet dispensing after the reset of a new patch.

The reward interval schedule for the depleting patch was calculated using the following equation:

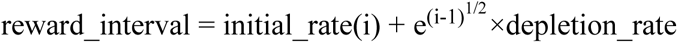

where *i* represents the sequence number of sugar pellets after a new patch reset.

### QUANTIFICATION AND STATISTICAL ANALYSIS

All statistical analyses were conducted using custom scripts implemented in MATLAB 2022b. Comparisons between two groups were performed using Welch’s t-test across all trials. For analyses involving two-by-two comparisons with unequal sample sizes, we applied a two-way ANOVA, followed by post-hoc multiple comparisons using Tukey’s HSD to determine pairwise differences. Time-series data were analyzed via repeated-measures MANOVA. Prior to applying parametric tests, data were evaluated for normality and homogeneity of variances. Error bars throughout represent the standard error of the mean (SEM).

### FORAGING EFFICIENCY CALCULATION

For each subject, foraging efficiency was calculated as the ratio of the total number of sugar pellets obtained to the product of the total distance moved and the subject’s body weight. Specifically, foraging efficiency was computed using the following equation:

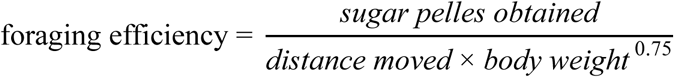

where “Sugar pellets obtained” is the count of reward pellets collected, “Distance moved” is the cumulative path length traversed by the subject during the foraging session, and “Body weight” is the mass of the subject measured prior to the session. This metric expresses the amount of reward obtained per unit of locomotor effort per unit body mass, thereby standardizing for individual differences in movement and size.

## Figure Legends

**Figure S1.**
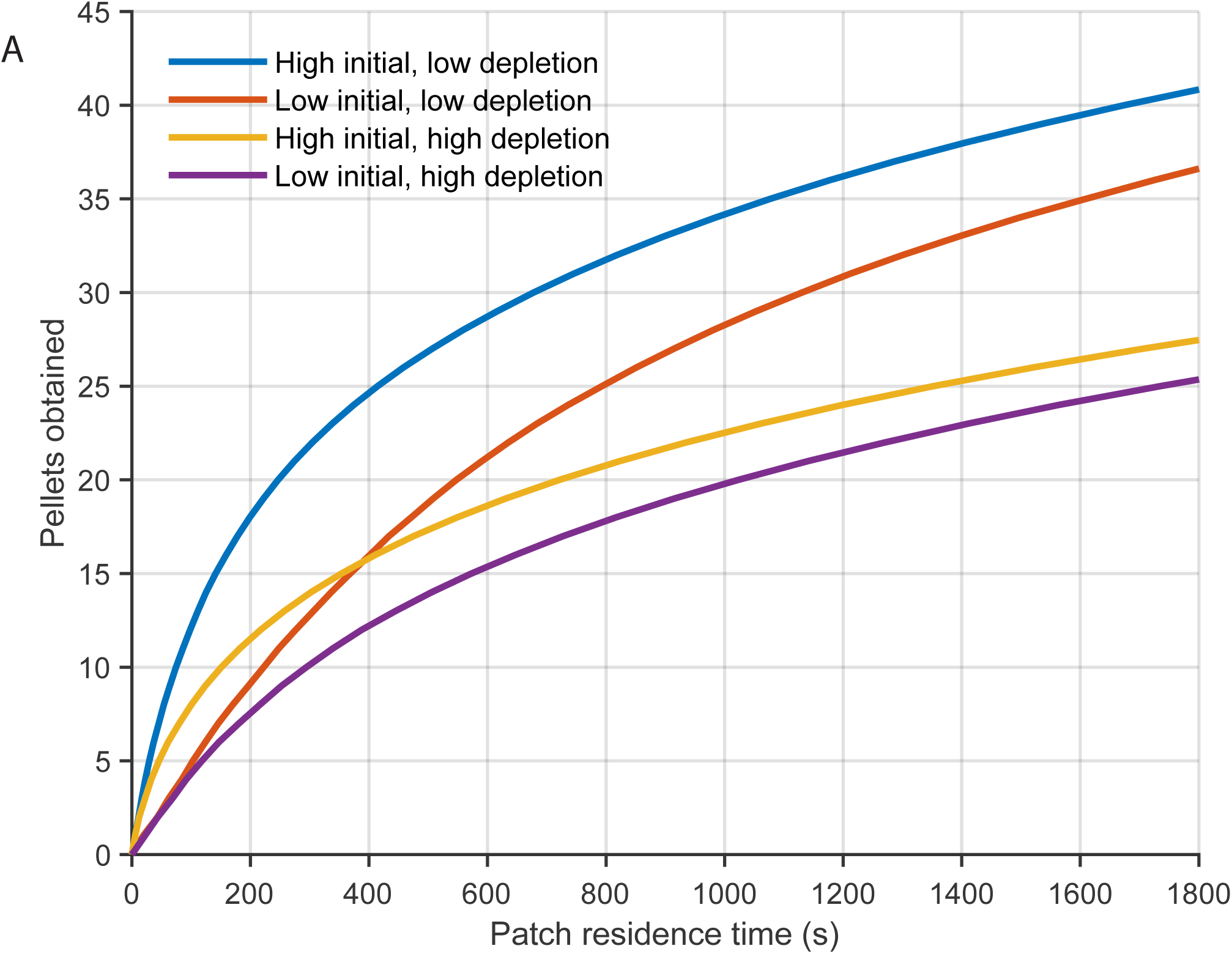
Pellet distribution functions used in the patch-leaving foraging task. (A) Schematic illustration of the four pellet distribution functions used in the task, defined by combinations of initial reward rate (high or low) and depletion rate (high or low). Pellet delivery rate decreased over time within each patch visit according to the specified depletion function.

